# Endosidin20 is a broad-spectrum cellulose synthesis inhibitor with an herbicidal function

**DOI:** 10.1101/2020.02.22.961094

**Authors:** Lei Huang, Chunhua Zhang

## Abstract

Cellulose is an important component of plant cell wall that controls anisotropic cell growth. Disruption of cellulose biosynthesis often leads to inhibited cell growth. Endosidin20 (ES20) was recently identified as a cellulose biosynthesis inhibitor (CBI) that targets the catalytic domain of Arabidopsis cellulose synthase 6 (CESA6) to inhibit plant growth. Here, we characterized the effects of ES20 on the growth of some other plant species and found that ES20 is a broad-spectrum plant growth inhibitor. We compared the inhibitory effects of ES20 and other CBIs on the growth of *cesa6* plants that have reduced sensitivity to ES20. We found that most of the *cesa6* with reduced sensitivity to ES20 show normal inhibited growth by other CBIs. ES20 also shows synergistic inhibitory effect on plant growth when applied together with other CBIs. We show ES20 has a different mode of action than tested CBIs isoxaben, indaziflam and C17. ES20 not only inhibits Arabidopsis growth under tissue culture condition, it inhibits plant growth under soil condition after direct spraying. We demonstrate that plants carrying two missense mutations can tolerate dual inhibition by ES20 and isoxaben.

**One sentence summary:** Cellulose biosynthesis inhibitor Endosidin20 has synergistic effect with other cellulose synthesis inhibitors and has the potential to be used as a spray herbicide.

## Introduction

Cellulose microfibril is crystalized polymer of β-1,4-D-glucose that serves as the main load-bearing component in plant cell wall. Cellulose is synthesized by rosette structured cellulose synthase complex (CSC) at the plasma membrane (PM) (Mueller et al., 1976; Giddings et al., 1980; Mueller and Brown, 1980; Pear et al., 1996; Arioli et al., 1998). Each CSC consists of 18 to 36 heterotrimeric cellulose synthases (CESAs) at 1:1:1 molar ratio (Doblin et al., 2002; Fernandes et al., 2011; Newman et al., 2013; Gonneau et al., 2014; Hill et al., 2014). The CSCs that synthesize the primary cell wall are composed of CESA1, CESA3 and CESA6 or CESA6-like subunit (CESA2, 5, or 9) whereas the CSCs synthesize the secondary cell wall are composed of CESA4, CESA7 and CESA8 (Taylor et al., 2003; Desprez et al., 2007; Persson et al., 2007). Rosette structured CSCs are located at the PM, Golgi and post-Golgi vesicles in electron microscope images of freeze-fractured plant cells (Haigler and Brown, 1986). Live cell imaging using functional fluorescence-tagged CESA also shows that CSCs are localized at the PM, Golgi, Trans-Golgi Network (TGN), and vesicles called microtubule-associated CESA compartments (MASCs) or small CESA compartments (SmaCCs) (Paredez et al., 2006; Crowell et al., 2009; Gutierrez et al., 2009). CSCs at the PM undergo bidirectional movement at the PM using microtubules as a guide and powered by cellulose polymerization (Paredez et al., 2006; Fujita et al., 2013). CSC subcellular transport requires the vesicle trafficking machinery and other proteins that interact with CESA. STELLO interacts with multiple CESAs to control efficient CSC exit of Golgi (Zhang et al., 2016), POM2/CELLULOSE SYNTHASE INTERACTIVE PROTEIN1(CSI) directly interacts with CESAs at the central cytoplasmic domain to associate CSCs with microtubules (Gu et al., 2010; Bringmann et al., 2012; Lei et al., 2012), and COMPANION OF CELLULOSE SYNTHASE1 (CC1) interacts with CESAs to regulate CSC transport under salt stress condition (Endler et al., 2015). Successful CSC delivery to the PM also requires the coordinated functions of actin, myosin XI, exocyst complex and PATROL1 (PTL1) (Sampathkumar et al., 2013; Zhu et al., 2018; Zhang et al., 2019). Newly identified SHOU4 protein negatively regulates CSC delivery to the PM and Clathrin-mediated endocytosis removes CSC from the PM (Bashline et al., 2013; Polko et al., 2018). Thus, cellulose biosynthesis is a complex process requires coordinated function of multiple proteins.

Cellulose biosynthesis inhibitors (CBIs) are small molecules that inhibit cellulose biosynthesis by targeting CESAs or other proteins required for cellulose synthesis. The CBIs often inhibit plant growth, cause cell swollen, and/or affect CSC subcellular localization (DeBolt et al., 2007; Harris et al., 2012; Brabham et al., 2014; Xia et al., 2014; Worden et al., 2015; Hu et al., 2016; Tateno et al., 2016). Isoxaben is one of the well characterized CBIs that has been widely used in understanding the mechanisms of cellulose biosynthesis. Isoxaben was originally used as an herbicide to control broad-leaf weeds a few decades ago because of its high efficiency in inhibiting plant growth (Huggenberger and Gueguen, 1987; Jamet and Thoisydur, 1988; Brinkmeyer et al., 1989). It was later found that isoxaben affects plant cell wall composition (Heim et al., 1990) and single amino acid mutations in *CESA3* and *CESA6* genes led to resistance to isoxaben in plant growth (Scheible et al., 2001; Desprez et al., 2002), providing evidence that isoxaben inhibits plant growth by targeting CESAs. Live cell imaging of plants expressing fluorescence-tagged CESA treated with isoxaben shows that isoxaben treatment can rapidly deplete CSC from the PM (Paredez et al., 2006), which makes it a useful inhibitor in understanding the subcellular trafficking of CSCs. A recently characterized small molecule C17 also depletes CSCs from the PM and has inhibitory effects on plant cytokinesis, root growth, and cellulose biosynthesis (Hu et al., 2016). The mutations in CESA1, CESA3, and pentatricopeptide repeat (PPR)-like proteins can overcome the inhibitory effect of C17 on plant growth (Hu et al., 2016). Interestingly, the inhibitor of mitochondrial complex III can also reduce plants’ sensitivity to C17 treatment, indicating that C17 might have a complex mode of action instead of directly targets CESA to inhibit plant growth. C17 has inhibitory effect on broad plant species, indicating it might be a good candidate for herbicide development (Hu et al., 2019). Indaziflam is a potent CBI that has been commercialized as an herbicide (Brabham et al., 2014). Interestingly, indaziflam increases the abundance of CSC at the PM to inhibit cellulose biosynthesis with an unknown mechanism (Brabham et al., 2014). CESTRIN reduces cellulose content in plant cell wall and removes CSC from the PM, but its endogenous target protein has not been characterized (Worden et al., 2015). Morlin is an inhibitor of microtubule dynamics and in turn affects the trajectories of CSC at the PM (DeBolt et al., 2007). The collection of CBIs allows transient manipulation of cellulose biosynthesis process and provides candidate small molecules for herbicide development.

Endosidin20 (ES20) inhibits Arabidopsis cellulose biosynthesis by targeting the catalytic site of CESA6 (Huang et al., 2020). Multiple missense mutations in *CESA6* lead to reduced sensitivity to ES20 in plant growth (Huang et al., 2020). Here, we report the characterization of ES20 on its inhibitory effect on different plant species and compare the action of ES20 with isoxaben, indaziflam and C17. We show that ES20 is a broad-spectrum plant growth inhibitor with a different mode of action than isoxaben, indaziflam and C17. Most of the mutants that have reduced sensitivity to ES20 are sensitive to isoxaben, indaziflam and C17. ES20 also has synergistic effect with these tested CBIs in inhibiting plant growth. We show that ES20 has the potential to be used as a commercial herbicide and it is possible to create plants with reduced sensitivity to both ES20 and isoxaben by gene editing.

## Results

### ES20 is a broad-spectrum plant growth inhibitor

Previous characterization of ES20 activity in Arabidopsis shows that it targets the catalytic site of CESA6 that is composed of highly conserved amino acids in CESAs (Huang et al., 2020). High conservation in amino acids at the catalytic site indicates that ES20 might be a broad-spectrum plant growth inhibitor that targets CESAs in different plants. We first tested the effects of ES20 on different dicotyledon and monocotyledon plant species in their growth. We found that ES20 can significantly inhibit the root growth of dicotyledon plants dandelion (*Taraxacum officinale*), tobacco (*Nicotiana benthamiana*), tomato (*Solanum lycopersicum*), and soybean (*Glycine max*) at the concentration of 5 μM (Figure 1A-1H). ES20 also inhibits the growth of monocotyledon plants rice (*Oryza sativa*) and maize (*Zea mays*) at the concentration of 20 μM (Figure 1I-1L). ES20 inhibition on the growth of two common grass weeds Perennial Ryegrass (*Lolium perenne*) and Kentucky Bluegrass (*Poa pratensis*) requires a concentration of 50 μM (Figure 1M-1P). Among the plants we have tested, dandelion, Perennial Ryegrass, Kentucky Bluegrass, and previously tested Arabidopsis, are common weeds found in agricultural field and lawn. The inhibitory effects of ES20 on both dicotyledon and monocotyledon plants indicate that ES20 is a broad-spectrum plant growth inhibitor.

**Figure 1.**
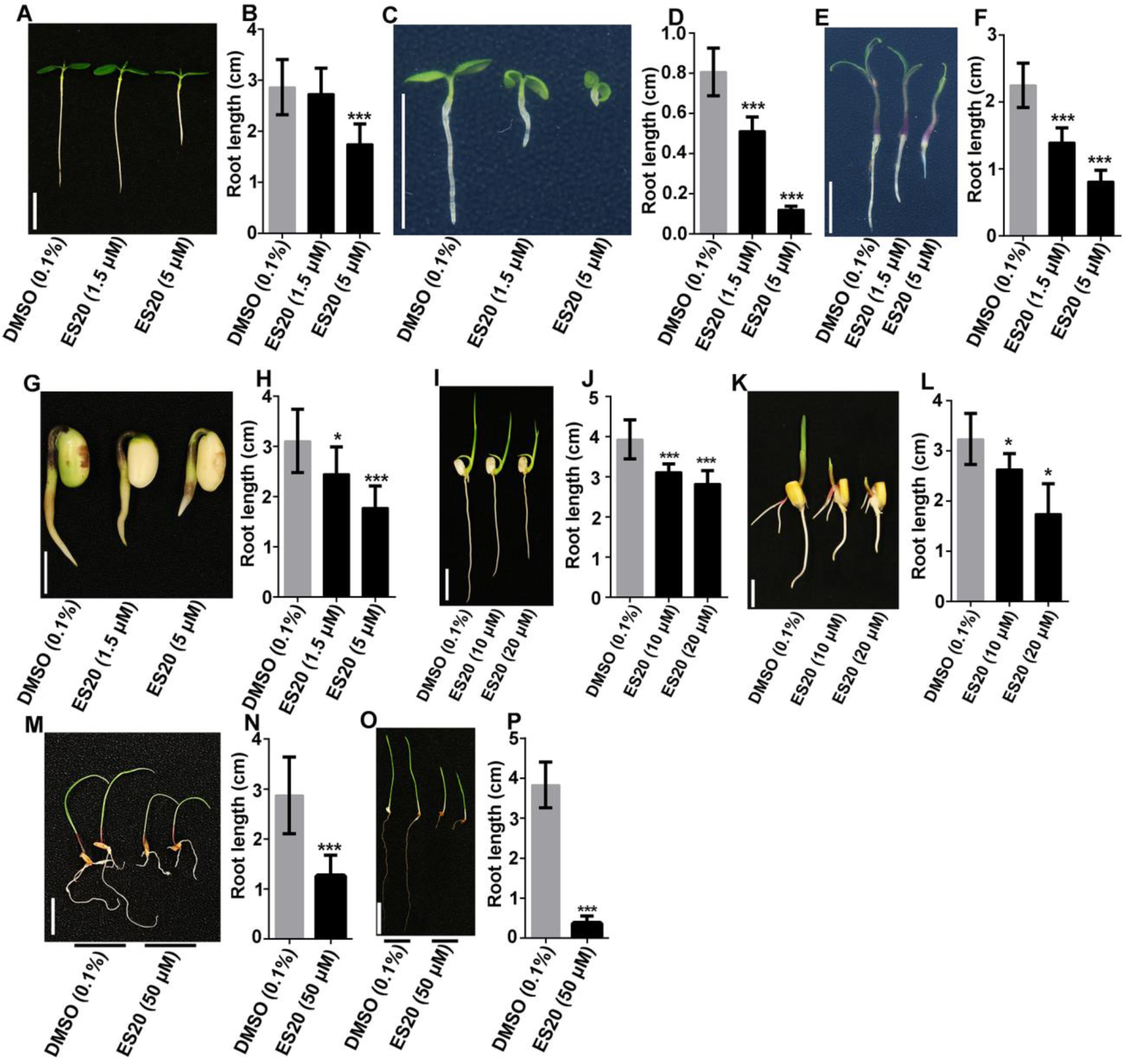
ES20 is a broad-spectrum plant growth inhibitor. A. representative seedlings of 5 days old dandelion (A), tobacco (C), tomato (E), soybean (G), rice (I), maize (K), perennial ryegrass (M) and Kentucky bluegrass (O) treated with DMSO (0.1%) and indicated concentration of ES20. Perennial ryegrass and Kentucky bluegrass seeds were soaked in sterile water supplemented with DMSO (0.1%) or ES20 (50 μM), whereas other plants’ seeds were grown on solid ½ MS growth medium supplemented with DMSO (0.1%) or indicated concentration of ES20. Scale bars: 1 cm. B, D, F, H, J, L, N and P, quantification of root length of A, C, E, G, I, K, M and O, respectively. * indicates p < 0.05 and *** indicates p < 0.001, by two-tailed student’s *t test* in comparison with DMSO treatment. Data represent mean ± SD. n= 10, 15, 10, 8, 12, 10, 10, 9 for panels B, D, F, H, J, L, N and P, respectively.

### Structure activity relationship analysis of ES20

ES20 (4-Methoxy-N-{[2-(2-Methylbenzoyl)Hydrazino]Carbothioyl}Benzamide) is a carbonothioyl benzamide derivative. In order to better understand the pharmacophore of ES20 that is essential for the inhibition of plant growth, we tested 11 ES20 analogs on plant growth (Figure 2A). We grew Arabidopsis Col-0 seedlings on 1 μM of different analogs and compared their root length with those of DMSO control (Figure 2B and 2C). Among the compounds we tested, only ES20 could significantly inhibit the root growth of Col-0, whereas none of the 11 analogs affected the root growth. After comparing the structures of the 11 analogs with that of ES20, we found that the 4-methoxy group and the position of this group is essential for ES20 inhibitory effect. The analogs that change 4-methoxy group to another group or change its position will not be active in inhibiting root growth. The methylbenzoyl group is also essential for ES20 activity. The analogs that alter the methylbenzoyl group by replacing the benzyl or change the position of the methyl will not be active in inhibiting plant growth.

**Figure 2.**
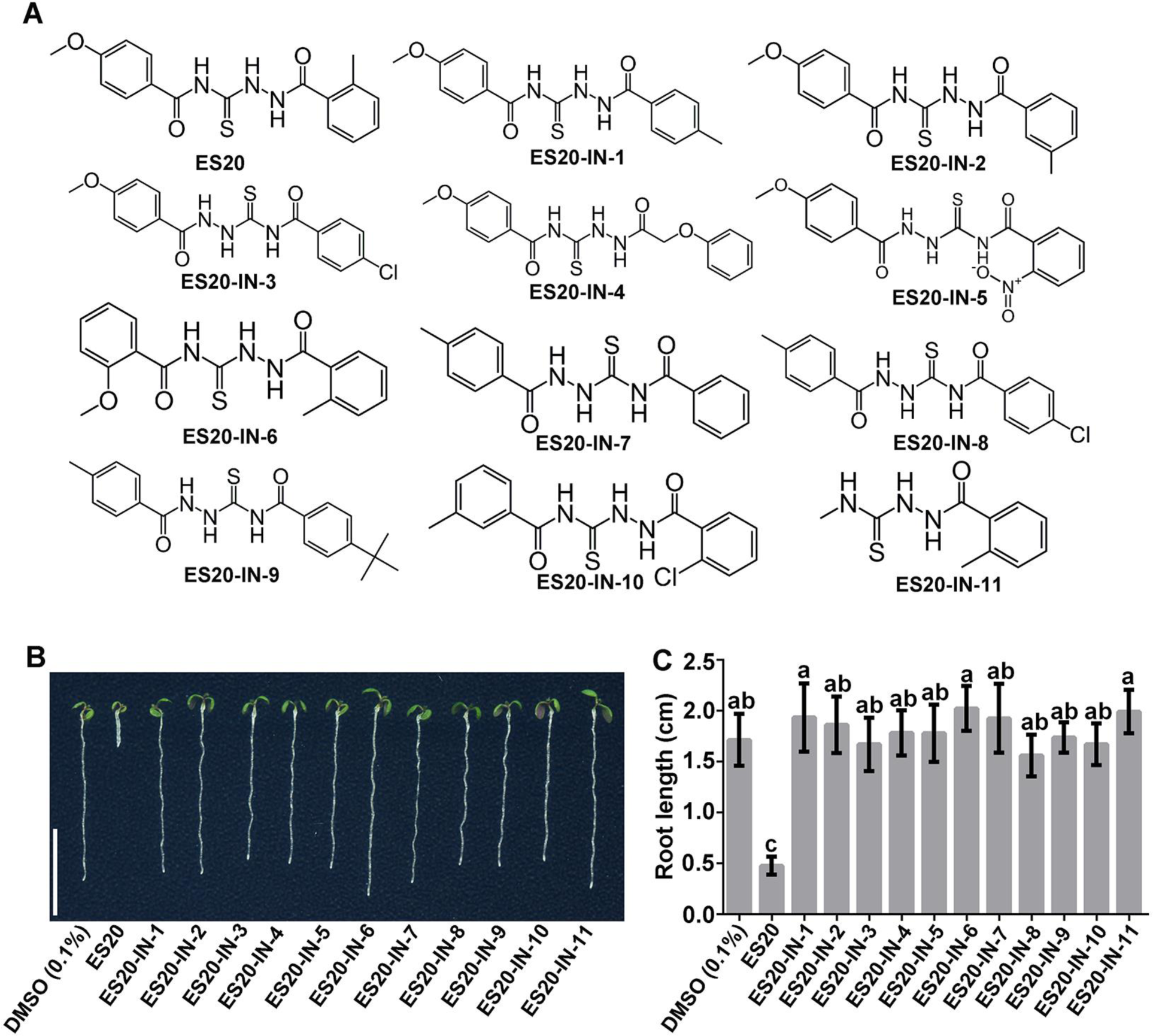
Structure activity relationship analysis of ES20. A. Chemical structures of ES20 and 11 analogs. B. 5 days old representative Arabidopsis Col-0 seedlings grew on ½ MS growth medium supplemented with DMSO (0.1%) or 1 μM different analogs. Scale bar: 1 cm. C. Quantification of root length of seedlings shown in B. Data represent mean ± SD. n = 10. The letters indicate statistically significant differences determined by one-way ANOVA tests followed by Tukey’s multiple comparison tests in different samples. Different letters indicate significant differences between groups (p < 0.05).

### ES20 uses a different mode of action than isoxaben, indaziflam and C17 to inhibit cellulose biosynthesis

The chemical structures of ES20, isoxaben, indaziflam and C17 are quite different (Supp. Figure S1), indicating they might use different modes of action to inhibit cellulose biosynthesis. Since the direct target protein for isoxaben, indaziflam, and C17 are not characterized, we compared their activities with that of ES20. We first tested whether our mutants that have reduced sensitivity to ES20 also have altered sensitivity to isoxaben, indaziflam and C17. We found that ES20, isoxaben, indaziflam and C17 can inhibit Arabidopsis growth at different efficiencies. Indaziflam is the most efficient in inhibiting plant growth. The root length of 5 days old Arabidopsis plants grown in the presence of 0.25 nM of indaziflam is only about 30% of those grown in the DMSO control media (Figure 3, SYP61-CFP and PIN2-GFP control plants). Isoxaben and C17 can inhibit Arabidopsis root growth for more than 50% at the concentration of 8 nM and 200 nM, respectively (Figure 3, SYP61-CFP and PIN2-GFP control plants). As reported previously, ES20 can inhibit about 80% of Arabidopsis root growth at the concentration of 1 μM (Figure 3, SYP61-CFP and PIN2-GFP control plants). When we grow *es20r* plants in growth media supplemented with 1 μM ES20, all of them show reduced sensitivity to ES20 when compared with the SYP61-CFP and PIN2-GFP control plants (Figure 3). When we grow these *es20r* mutants in the growth media supplemented with 8 nM isoxaben, 0.25 nM indaziflam, or 200 nM C17, most of the *es20r* plants display inhibited growth by these CBIs at a level similar to the control plants (Figure 3). However, we did notice some of the *es20r*s show reduced sensitivity to isoxaben, indaziflam and C17 (Figure 3). After quantification of the relative root growth inhibition, we find *esr20r1*, *esr20r3*, *esr20r4*, *esr20r5* and *esr20r10* have reduced sensitivity to isoxaben, *esr20r1*, *esr20r3*, *esr20r4*, *esr20r5* show reduced sensitivity to indaziflam, and *esr20r3*, *esr20r4*, *esr20r5*, *esr20r6*, *esr20r7* and *esr20r12* show reduced sensitivity to C17 (Figure 3). The mutants’ different sensitivity to these CBIs imply that ES20 has a different target site than other three CBIs.

**Figure 3.**
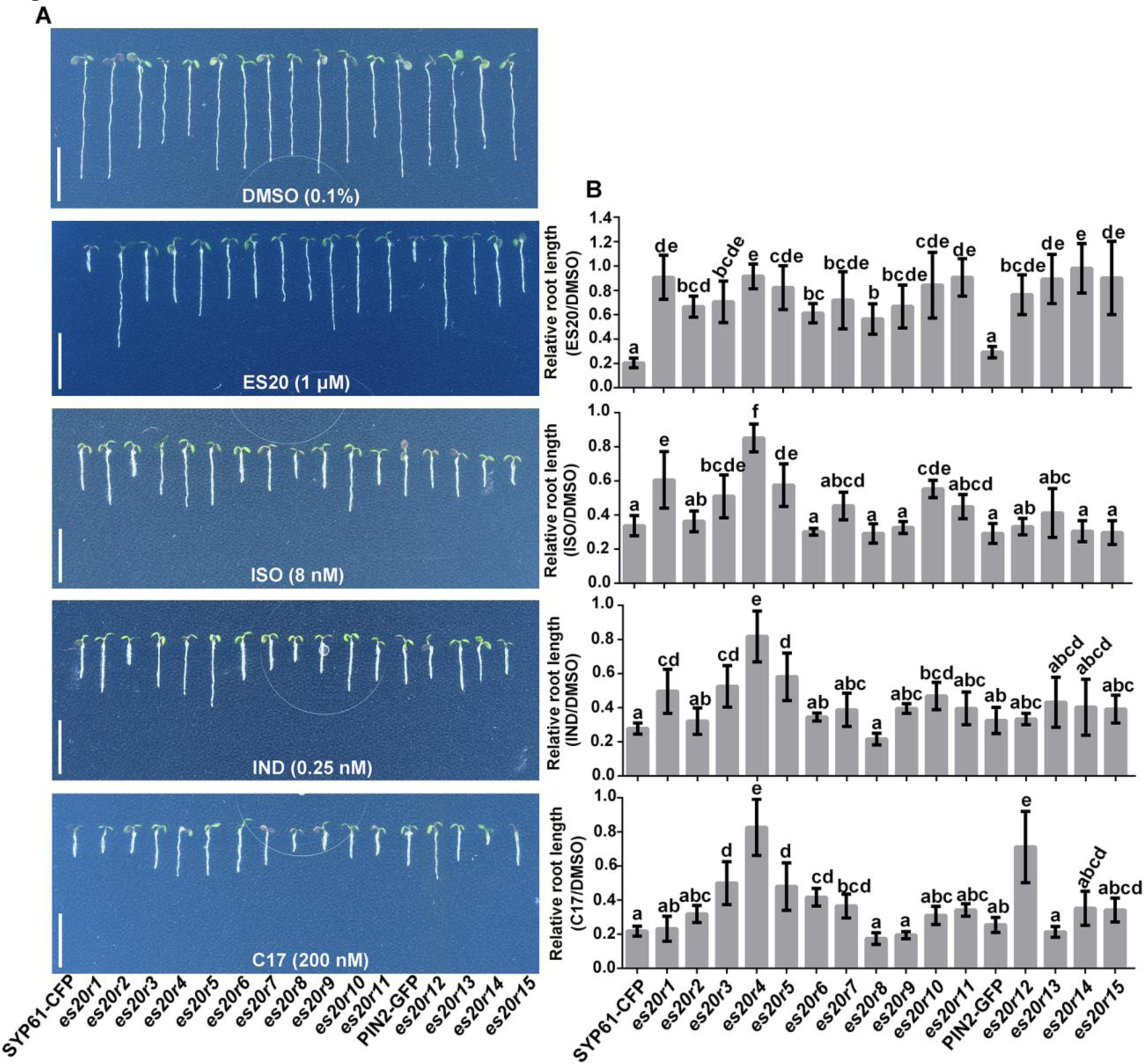
The growth of *es20r*s in the presence of isoxaben, indaziflam and C17. A. Representative seedlings of 5 days old *es20r*s grown on ½ MS medium supplemented with DMSO (0.1%), ES20 (1 μM), isoxaben (8 nM), indaziflam (0.25 nM) or C17 (200 nM). Scale bars: 1 cm. B. Quantification of the relative root length of seedlings as shown in A. The letters indicate statistically significant differences determined by one-way ANOVA tests followed by Tukey’s multiple comparison tests in different samples. Different letters indicate significant differences between groups (p < 0.05). Data represent mean ± SD. n = 12. ISO: isoxaben. IND: indaziflam.

In our previous study, we found that six predicted mutations at CESA6 catalytic site (D562N, D564N, D785N, Q823E, R826A, W827A) from modeled structure cause plants to have reduced sensitivity to ES20 in growth (Huang et al., 2020). To further test whether ES20 and other CBIs have the same binding site, we examined how the six predicted mutations at the catalytic site and two predicted mutations beyond the catalytic site (L365F and D395N) in CESA6 affect plants’ response to the three CBIs. Consistent with previous result, six predicted mutations at the catalytic site cause reduced sensitivity to ES20 and two predicted mutations beyond the catalytic do not affect plants’ sensitivity to ES20 (Figure 4), whereas none of the predicted mutations affects plants’ sensitivity to other three CBIs (Figure 4). Plants’ reduced sensitivity to ES20 caused by predicted mutations but normal sensitivity to other three CBIs further implies that ES20 has a different target site than other three CBIs.

**Figure 4.**
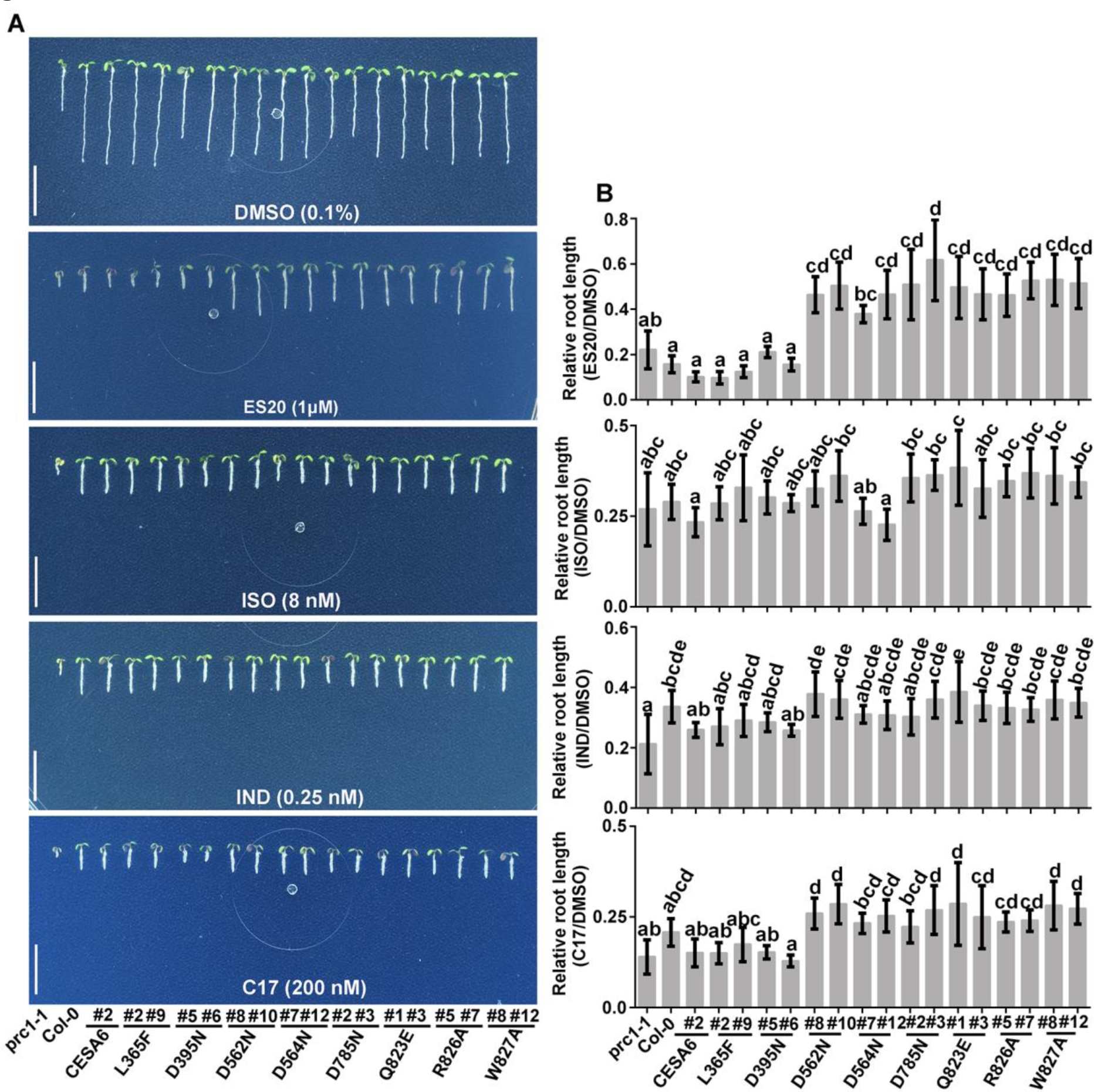
Plants expressing CESA6 carrying predicted mutations at the modeled binding site are sensitive to isoxaben, indaziflam and C17. A. 5 days old representative seedlings of *prc1-1/cesa6* complemented with wild type CESA6 or mutated CESA6 carrying predicted mutations at modeled catalytic site grown on ½ MS medium supplemented with DMSO (0.1%), ES20 (1 μM), isoxaben (8 nM), indaziflam (0.25 nM) or C17 (200 nM). Scale bars: 1 cm. B. Quantification of the relative root length of seedlings as mentioned in A. The letters indicate statistically significant differences determined by one-way ANOVA tests followed by Tukey’s multiple comparison tests in different samples. Different letters indicate significant differences between groups (p < 0.05). Data represent mean ± SD. n = 12. ISO: isoxaben. IND: indaziflam.

Three mutations, CESA3^G998D^ (*ixr1-1*), CESA3^T942I^ (*ixr1-2*) and CESA6^R1064W^ (*ixr2-1*) have been found to cause reduced sensitivity to isoxaben (Scheible et al., 2001; Desprez et al., 2002). Isoxaben is thus believed to target CESA directly to inhibit cellulose biosynthesis and has been widely used in cellulose biosynthesis research (Scheible et al., 2001; Desprez et al., 2002; Shim et al., 2018). The originally identified mutations lead to reduced sensitivity to isoxaben are located at the C-terminal region of CESAs while most of the mutations that cause reduced sensitivity to ES20 are located at the central cytoplasmic domain. We tested how the isoxaben insensitive mutants respond to ES20. We grew *ixr1-1*, *ixr1-2* and *ixr2-1* on growth medium supplemented with DMSO (0.1%), isoxaben (10 nM) or ES20 (1 μM) for five days. We found the three *ixr* mutants display reduced sensitivity to isoxaben but these mutant plants have the same sensitivity to ES20 as wild type plants (Figure 5). The normal response of isoxaben resistant plants to ES20 also indicates ES20 targets CESA differently than isoxaben.

**Figure 5.**
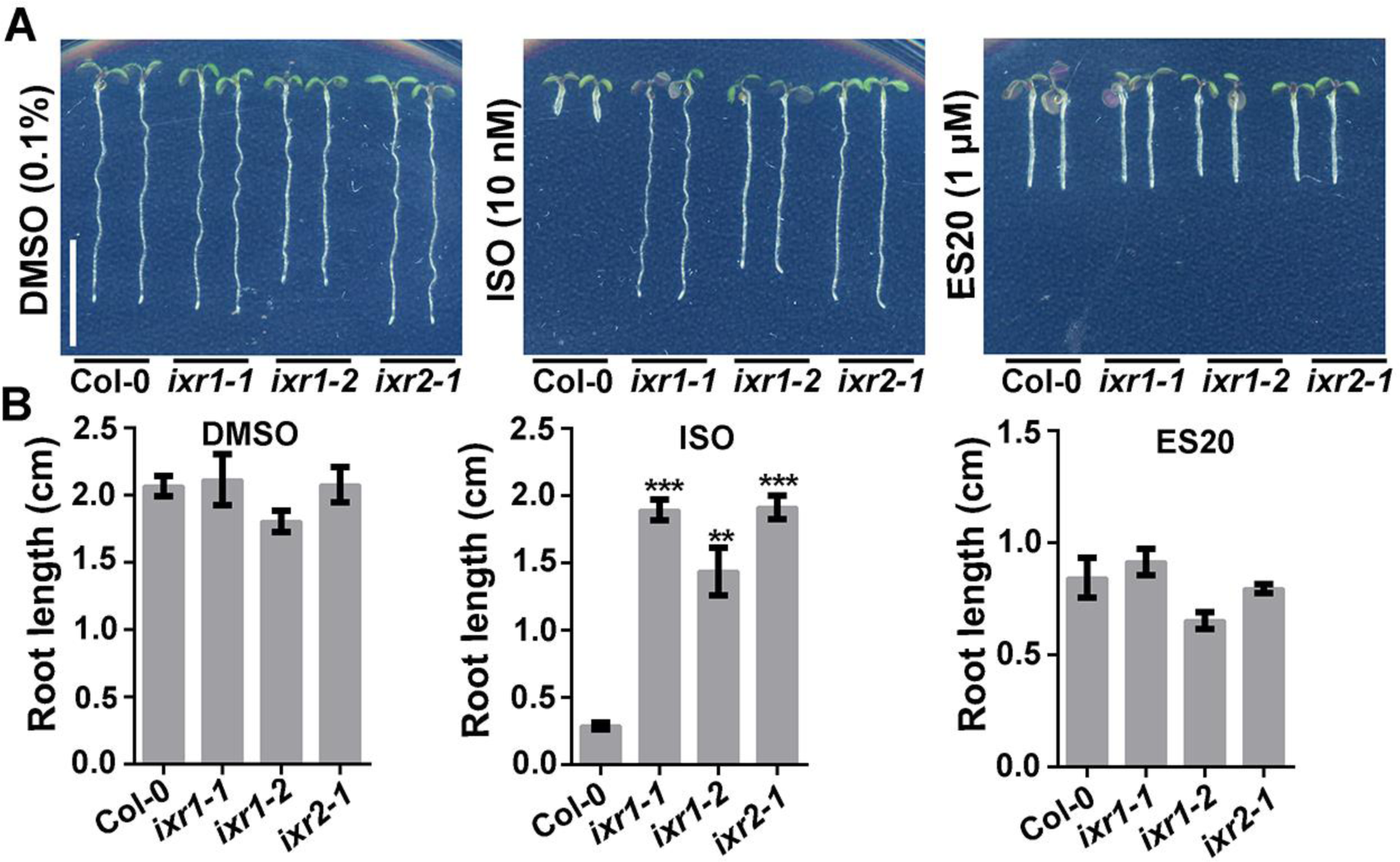
Isoxaben insensitive mutants are sensitive to ES20. A. Representative seedlings of 5 days old Col-0 and three isoxaben insensitive mutants (*ixr1-1*, *ixr1-2* and *ixr2-1*) grown on ½ MS growth medium supplemented with DMSO (0.1%), isoxaben (10 nM) or ES20 (1 μM). Scale bar: 1 cm. B. Quantification of root length of seedlings as shown in A. ** indicates p < 0.01, *** indicates p < 0.001, by two-tailed student’s t test in comparison with Col-0. Data represent mean ± SD. n= 9. ISO: isoxaben.

### ES20 has synergistic inhibition effect on plant growth with other CBIs

Since ES20 has a different mode of action compared with isoxaben, indaziflam and C17, we wonder whether ES20 also has synergistic effects with these CBIs in inhibiting plant growth. We first did a series of concentration test for ES20, isoxaben, indaziflam and C17 to determine the maximum concentration for each that will not inhibit the root growth of Col-0 seedlings. As shown in Figure 6 and Figure S2, 250 nM ES20, 4 nM isoxaben, 0.06 nM indaziflam or 40 nM C17 alone does not significantly inhibit wildtype plant root growth. However, when we did the dual drug treatments, we found that 250 nM ES20 mixed with 4 nM isoxaben, 0.06 nM indaziflam or 40 nM C17 could significantly inhibit the root growth compared with DMSO control treatment or the single drug treatment (Figure 6, Figure S2). Synergistic effects of ES20 with other CBIs in inhibiting root growth further indicates that ES20 has a different mode of action than isoxaben, indaziflam and C17.

**Figure 6.**
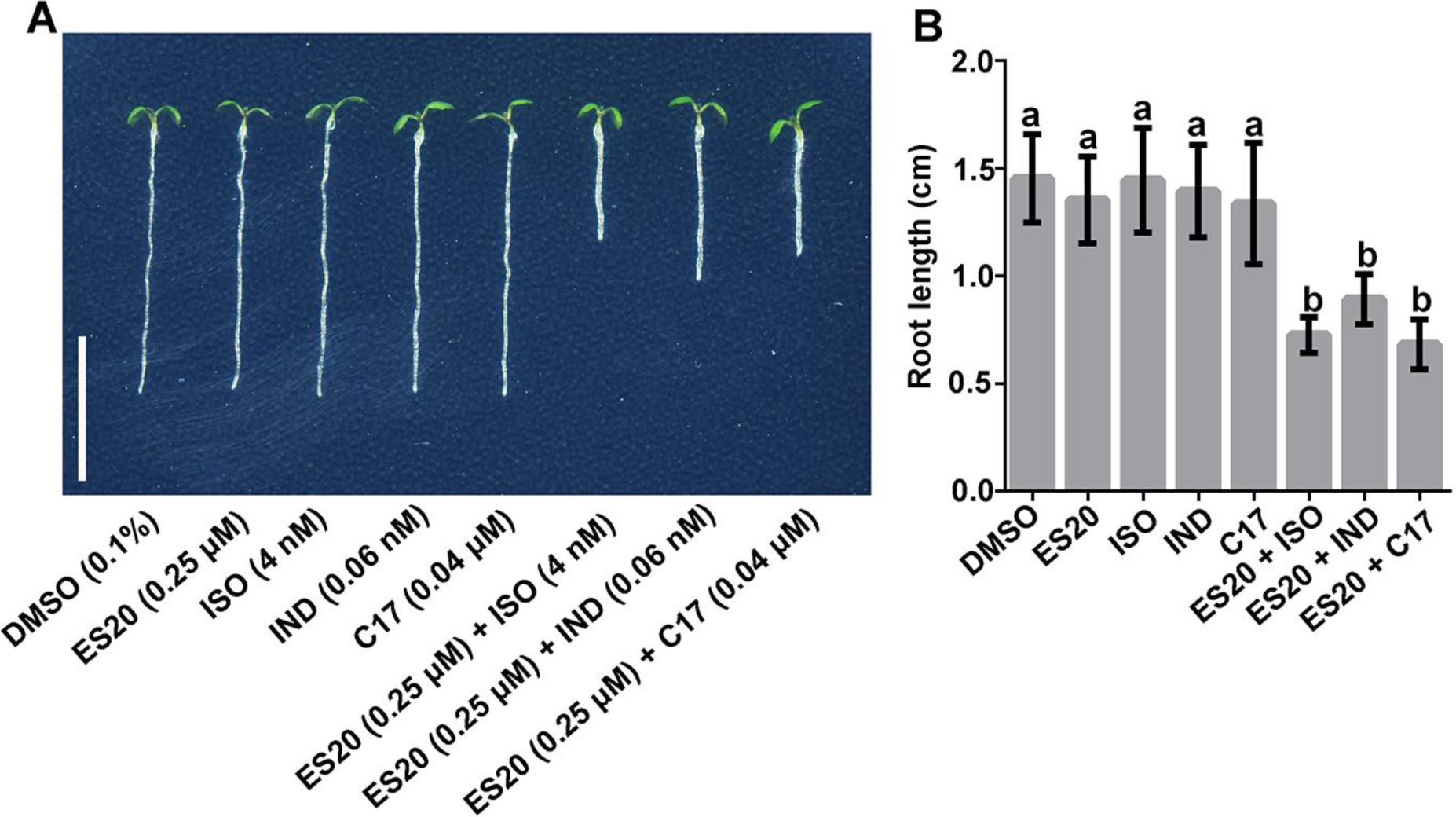
ES20 has synergistic inhibition effect on root growth with isoxaben, indaziflam and C17. A. Representative seedlings of 5 days old Col-0 grown on ½ MS medium supplemented with DMSO (0.1%), ES20 (0.25 μM), isoxaben (4 nM), indaziflam (0.06 nM), C17 (0.04 μM) and a mixture of ES20 (0.25 μM) with three other inhibitors. Scale bar: 1 cm. B. Quantification on the root length of seedlings as shown in A. The letters indicate statistically significant differences determined by one-way ANOVA tests followed by Tukey’s multiple comparison tests in different samples. Different letters indicate significant differences between groups (p < 0.05). Data represent mean ± SD. n = 15. ISO: isoxaben. IND: indaziflam.

### Editing on *CESA6* allows plants to tolerate ES20 inhibition without affecting growth

Previous chemical genetic screens allow us to obtain 15 *CESA6* mutants that have reduced sensitivity to ES20 in growth. Among these mutants, *es20r1* (CESA6^E929K^) does not have significantly reduced root growth by itself and displays least level of growth inhibition by ES20 (Figure 2) (Huang et al., 2020). Normal growth and strong tolerance to ES20 make it a promising approach to edit *CESA6* to create ES20-tolerant plants. We introduced a single nucleotide mutation in YFP-CESA6 genomic construct to create YFP-CESA6^E929K^ construct. We then transformed YFP-CESA6 and YFP-CESA6^E929K^ constructs to *cesa6* null mutant *prc1-1*. We then screened for single insertion lines for both YFP-CESA6 and YFP-CESA6^E929K^ and obtained independent homozygous single insertion transgenic lines for YFP-CESA6 and YFP-CESA6^E929K^. We found that expression of YFP-CESA6 can rescue the growth defect of *prc1-1* and the transgenic plants have normal sensitivity to ES20 inhibition when the plants are grown on growth medium supplemented with ES20 (Figure 7A and 7B). However, YFP-CESA6^E929K^ can not only rescue the growth defect of *prc1-1*, the transgenic plants display tolerance to ES20 inhibition in root growth when grown on growth media supplemented with ES20 (Figure 7A and 7B). We also grew the transgenic plants on normal growth media and then treated the seedlings with ES20 overnight. We found that YFP-CESA6;*prc1-1* plants are swollen and have increased root diameter at root tips after ES20 treatment (Figure 7C and 7D). However, YFP-CESA6^E929K^;*prc1-1* plants are not swollen under the same ES20 treatment condition (Figure 7C and 7D). The growth assays indicate that CESA6^E929K^ mutation is sufficient in causing plants to tolerate ES20 inhibition in growth.

**Figure 7.**
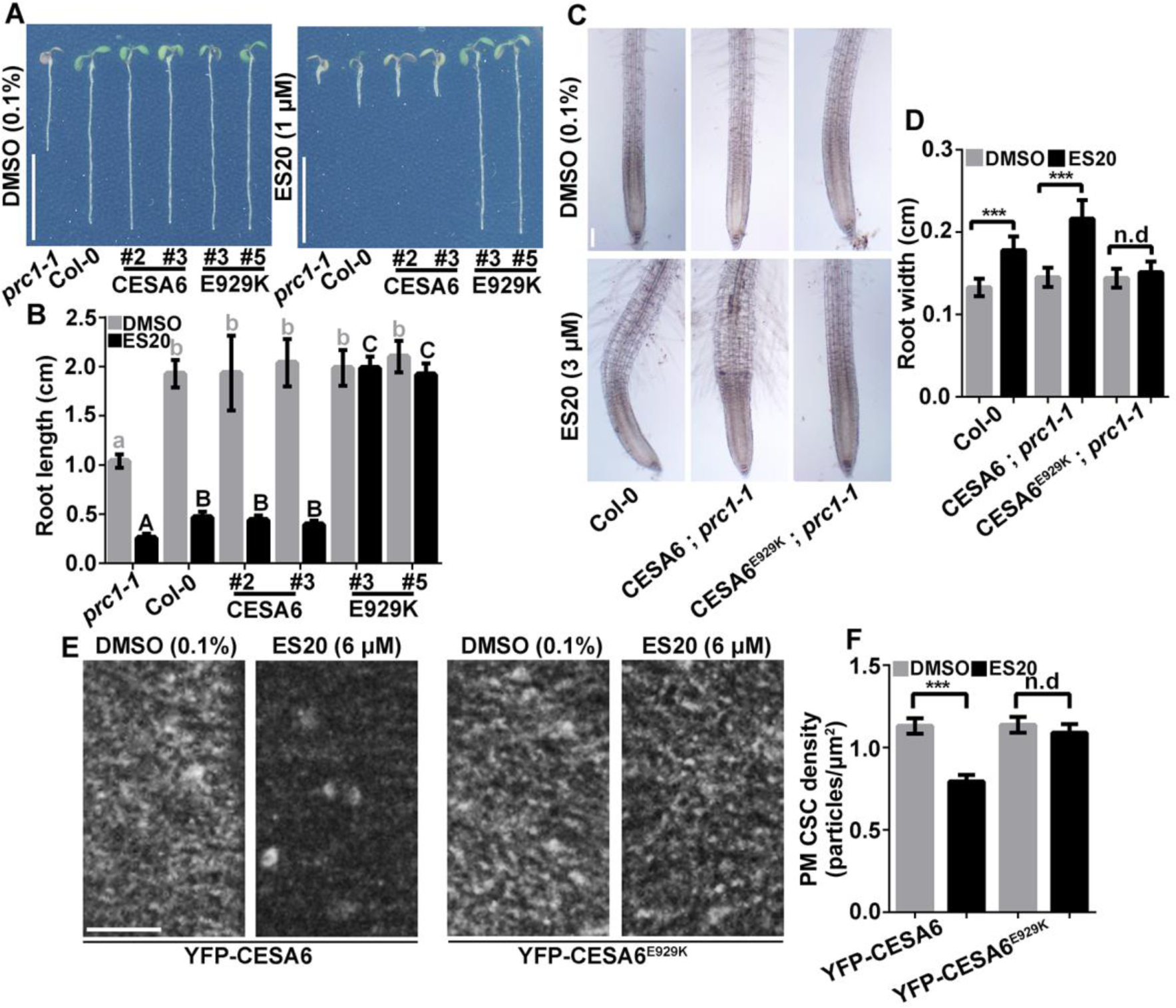
CESA6 point mutation E929K abolishes ES20’s inhibitory effect on root growth and on the depletion of CSC localization at the PM. A. Representative seedlings of 5 days old *prc1-1*, Col-0 and *prc1-1* complemented with wild type or mutated CESA6 constructs grown on the ½ MS medium supplemented with DMSO (0.1%) or ES20 (1 μM). Scale bars: 1 cm. B. Quantification on the root length of seedlings as shown in A. The letters indicate statistically significant differences determined by one-way ANOVA tests followed by Tukey’s multiple comparison tests in different samples. Different letters indicate significant differences between groups (p < 0.05). Lower- and upper-case letters represent ANOVA analysis of plants grown on media with DMSO and ES20, respectively. Data represent mean ± SD. n = 10. C and D. CESA6 point mutation E929K abolishes cell swollen phenotype caused by ES20 treatment. C. Representative root images of 5 days old Col-0 and transgenic plants expressing wildtype or mutated CESA6 in *prc1-1* background treated with liquid ½ MS supplemented with DMSO (0.1%) or ES20 (3 μM) for 20 hours. Scale bars: 100 μm. D. Quantification on the root width of seedlings as shown in C. *** indicates p < 0.001, by two-tailed student’s t test in comparison with DMSO treatment, while n.d indicates no significant difference. Data represent mean ± SD. n = 15. E and F. The mutation E929K causes reduced sensitivity to the effect of ES20 treatment on CSC localization. E. Representative images of PM-localized YFP-CESA6 and YFP-CESA6^E929K^ after 30 min ES20 treatment. Scale bar: 5 μm. F. Quantification on the density of PM localized CSC as shown in E. *** indicates p < 0.001, by two-tailed student’s t test in comparison with DMSO treatment, while n.d indicates no significant difference. Data represent mean ± SE. n = 24.

ES20 targets CESA6 and short-term ES20 treatment causes reduced CSC localization at the PM (Huang et al., 2020). Since YFP-CESA6^E929K^ is sufficient in causing plants to be tolerant to the growth inhibition and cell swollen caused by ES20, we wonder whether the tolerance occurs at the cellular level as well. We performed short-term ES20 treatment on YFP-CESA6;*prc1-1* and YFP-CESA6^E929K^;*prc1-1* plants and examined CSC localization. Consistent with previous report, YFP-CESA6;*prc1-1* seedlings treated with 6 μM ES20 for 30 min have significantly reduced CSC density at the PM compared with the DMSO control treatment (Figure 7E and 7F). However, 30 min of 6 μM ES20 treatment does not significantly affect the CSC density at the PM in CESA6^E929K^;*prc1-1* seedlings (Figure 7E and 7F). Thus, a single amino acid change in CESA6 is sufficient to cause plants to tolerate ES20 in plant growth and in CSC trafficking at the cellular level.

Our previously identified *cesa6* alleles with reduced sensitivity to ES20 provide guidance for generating other plant species with reduced sensitivity to ES20 through genetic engineering method. In order to test whether the reduced ES20 sensitivity trait is dominant or recessive, we transformed three YFP-CESA6 genomic constructs with native *CESA6* promoter carrying missense mutations (YFP-CESA6^E929K^, YFP-CESA6^T783I^ and YFP-CESA6^D396N^) to Arabidopsis wildtype Col-0 through agrobacterium-mediated transformation. We grew transgenic plants expressing YFP-CESA6^E929K^, YFP-CESA6^T783I^ and YFP-CESA6^D396N^ on growth medium supplemented with DMSO (0.1%) or ES20 (1 μM). These transgenic plants do not have obvious growth defects compared with Col-0 when grown on DMSO control medium (Figure 8). However, the transgenic plants expressing YFP-CESA6^E929K^, YFP-CESA6^T783I^ and YFP-CESA6^D396N^ have longer roots than YFP-CESA6 plants when grown on growth media supplemented with ES20 (Figure 8). We also noticed that plants expressing mutated CESA6 constructs in Col-0 background have lower level of ES20 tolerance than those of the EMS mutants (Figure 3, Figure 8), indicating reduced sensitivity to ES20 caused by CESA6 mutations are semi-dominant.

**Figure 8.**
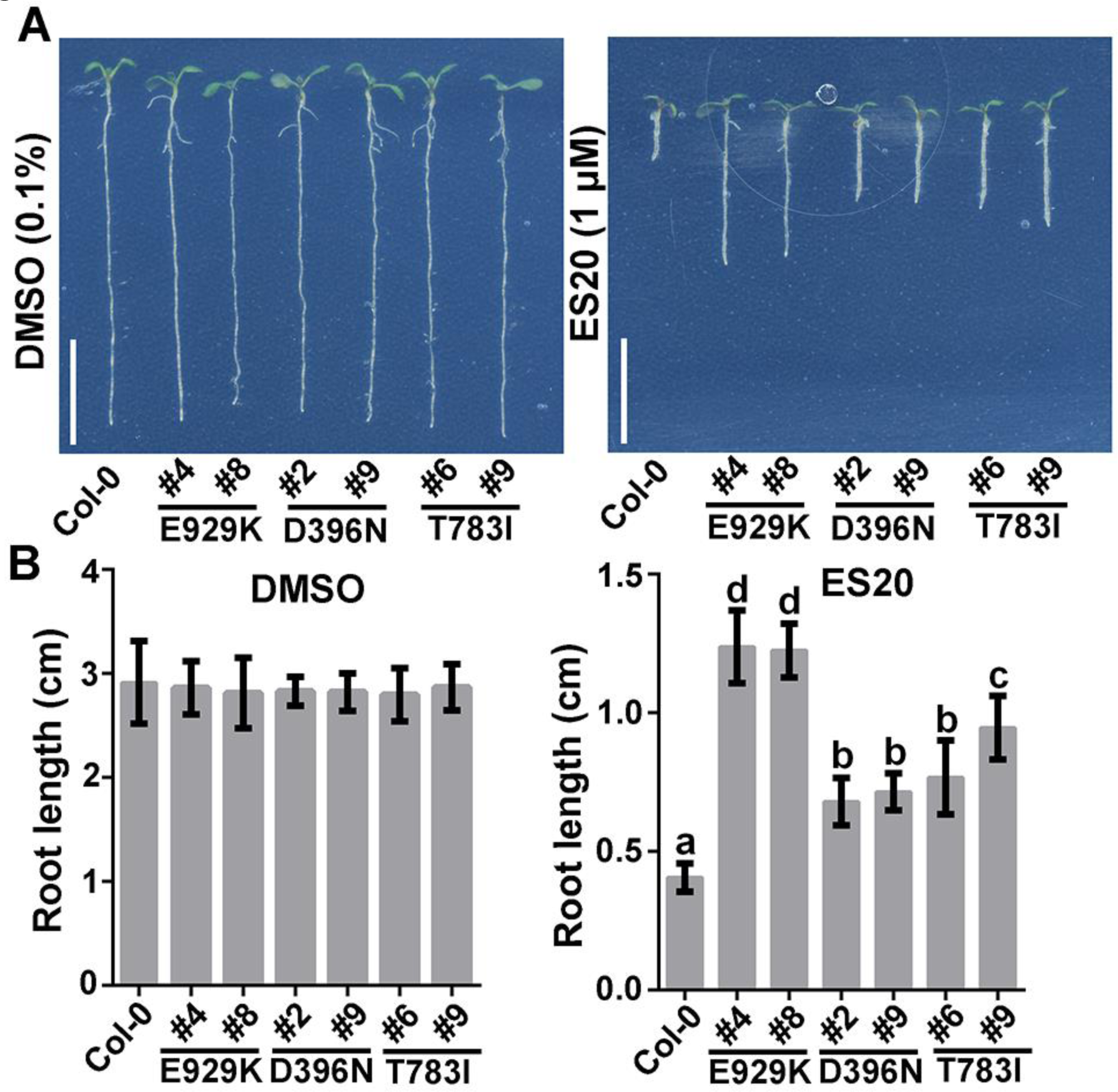
ES20 tolerance caused by CESA6 mutations is a semi-dominant trait. A. Representative seedlings of 5 days old Col-0 and transgenic lines expressing three different mutated CESA6 constructs (CESA6^E929K^, CESA6^D396N^ and CESA6^T783I^) in Col-0 grown on ½ MS medium supplemented with DMSO (0.1%) and ES20 (1 μM). Scale bars: 1 cm. B. Quantification on the root length of seedlings as shown in A. The letters indicate statistically significant differences determined by one-way ANOVA tests followed by Tukey’s multiple comparison tests in different samples. Different letters indicate significant differences between groups (p < 0.05). Data represent mean ± SD. n = 10. ISO: isoxaben.

### Generation of ES20 and isoxaben dual tolerant plant

Long time repetitive application of single herbicide could be problematic since herbicide tolerant weeds emerge by natural mutation due to the single selective pressure (Heap, 2014). Since ES20 and isoxaben seem to target CESA at different binding site, application of ES20 and isoxaben together is expected to reduce the chance of herbicide tolerant weed development. Establishing a strategy to create crop plants that are resistant to both ES20 and isoxaben is expected to be important for using ES20 and isoxaben for weed control in agricultural production. We tried to combine ES20 and isoxaben tolerant trait in plants by crossing the isoxaben insensitive mutant *ixr1-1* with *es20r1*. We obtained the homozygous *ixr1-1*;*es20r1* lines in F3 generation and tested the growth phenotype on growth medium supplemented with DMSO (0.1%), ES20 (1 μM), isoxaben (12 nM) and ES20 (1 μM) plus isoxaben (12 nM). As shown in Figure 9A and Figure 9B, *ixr1-1* and *es20r1* seedlings do not have obvious root growth defects compared with wild type plants when grown on growth media supplemented with DMSO. However, *ixr1-1*;*es20r1* double mutant plants have slightly reduced root length compared with wild type (Figure 9A and 9B). The single mutant plants of *ixr1-1* and *es20r1* have reduced sensitivity to isoxaben and ES20, respectively (Figure 9A and 9B). However, the *ixr1-1*;*es20r1* double mutants can tolerate the mixture of ES20 (1 μM) and isoxaben (12 nM) treatment (Figure 9A and 9B). As *ixr1-1*;*es20r1* double mutant plants have slightly reduced root growth at seedling age, we wanted to see whether there will be growth phenotype in later growth stage. We grew the mutant plants in the soil till the end of their life cycle and found that *ixr1-1* single mutant and *ixr1-1*;*es20r1* double mutant plants have smaller rosette when compared with wild type (Figure 9C and 9D). The height of 40 days old soil grown *ixr1-1*;*es20r1* double mutant plant is also shorter compared with wild type and the single mutants (Figure 9E and 9F). Thus, although the double mutant of *ixr1-1*;*es20r1* can tolerate both ES20 and isoxaben, there is some trade off in growth.

**Figure 9.**
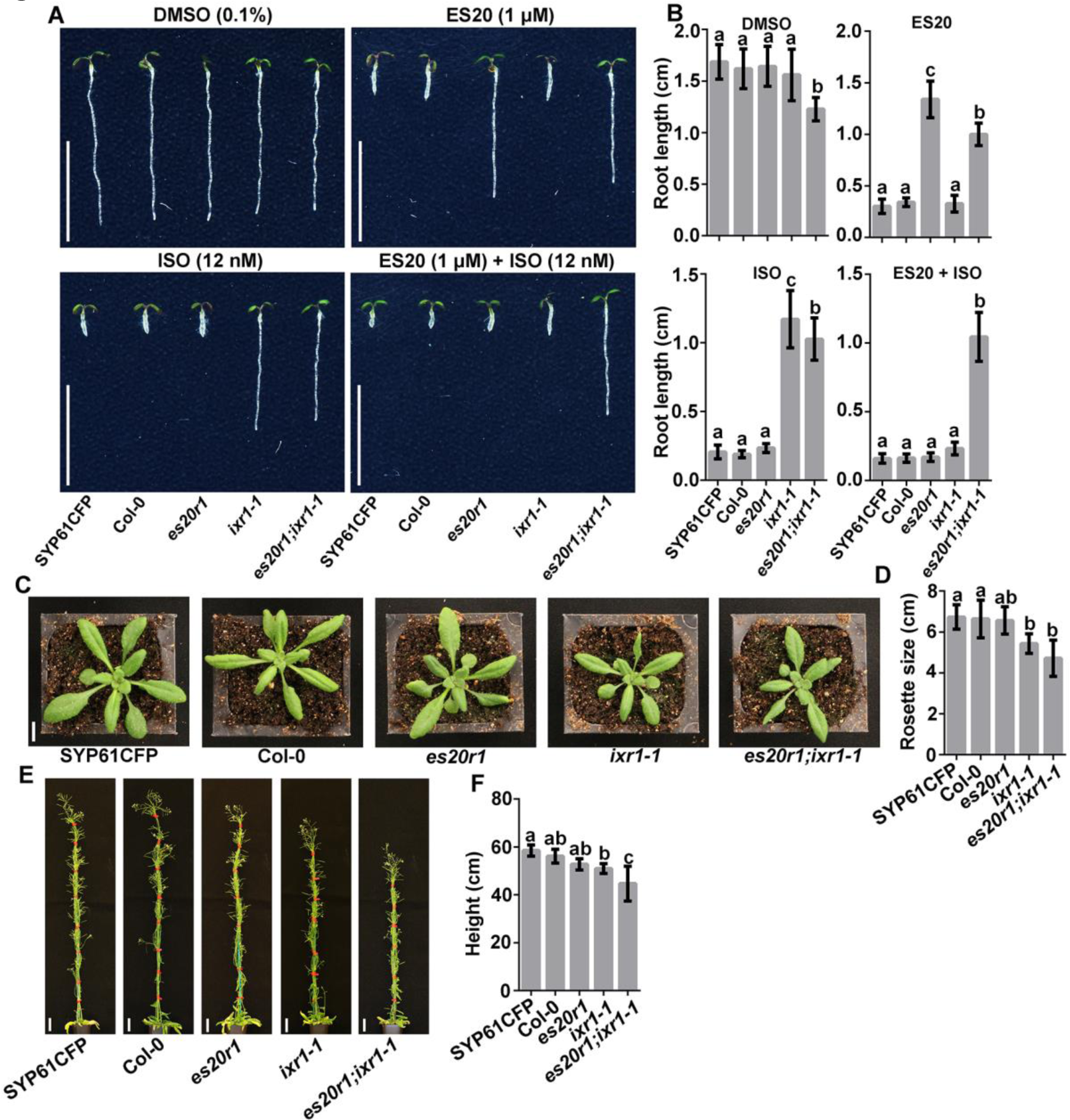
Double mutant *ixr1-1*;*esr20r1* can tolerate cotreatment of ES20 and isoxaben. A and B. *ixr1-1*;*es20r1* seedlings exhibits reduced sensitivity to the mixture of ES20 and isoxaben. A. Representative seedlings of 5 days old SYP61-CFP, Col-0, *es20r1*, *ixr1-1* and *es20r1*;*ixr1-1* grown on ½ MS medium supplemented with DMSO (0.1%), ES20 (1 μM), isoxaben (12 nM) and the mixture of ES20 (1 μM) and isoxaben (12 nM). Scale bars: 1 cm. B. Quantification on the root length of seedlings as shown in A. The letters indicate statistically significant differences determined by one-way ANOVA tests followed by Tukey’s multiple comparison tests in different samples. Different letters indicate significant differences between groups (p < 0.05). Data represent mean ± SD. n = 15. C. The rosettes of 3 weeks old SYP61-CFP, Col-0, *es20r1*, *ixr1-1* and *es20r1;ixr1-1* grown on soil. Scale bar: 1 cm. D. Quantification on the size of rosettes in 3 weeks old soil grown plants as shown in C. Rosette size was measured as the sum of the lengths of the longest leaf and second longest leaf. Data represent mean ± SD. n = 9. The letters indicate statistically significant differences determined by one-way ANOVA tests followed by Tukey’s multiple comparison tests in different samples. Different letters indicate significant differences between groups (p < 0.05). E. Representative plants of 40 days old soil grown SYP61-CFP, Col-0, *es20r1*, *ixr1-1* and *es20r1;ixr1-1.* Scale bars: 3 cm. F. Quantification on the height of 40 days soil grown plants as shown in E. Data represent mean ± SD. n = 8. The letters indicate statistically significant differences determined by one-way ANOVA tests followed by Tukey’s multiple comparison tests in different samples. Different letters indicate significant differences between groups (p < 0.05).

### ES20 has the potential to be used as a spray herbicide

To test whether ES20 could be used as a potential herbicide, we sprayed soil grown wildtype plant with ES20 to see whether it could inhibit the growth or even kill the soil grown plants after spraying. We transferred 5 days old Col-0 seedlings grown in growth medium to the soil and sprayed with 50 mL sterile water contained DMSO (0.5%) or ES20 (500 μM), respectively. 7 days after spraying, ES20 treated seedlings almost completely died while the DMSO treated seedlings showed normal growth (Figure 10). The small-scale spraying experiments indicate ES20 has the potential to be used as a spray herbicide.

**Figure 10.**
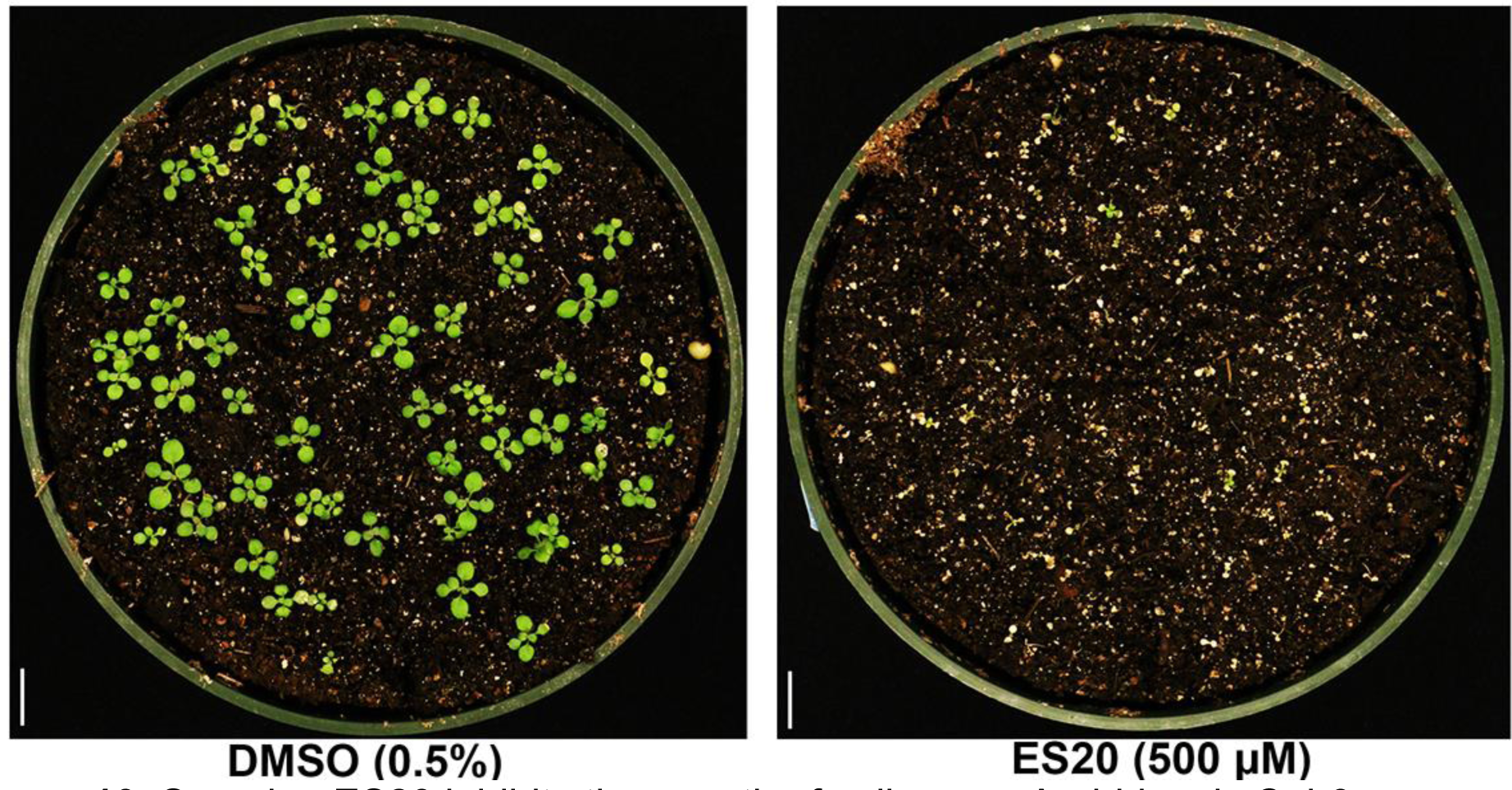
Spraying ES20 inhibits the growth of soil grown Arabidopsis Col-0. Arabidopsis Col-0 grown on soil sprayed with DMSO (0.5%) (left) and ES20 (500 μM) (right). Images were taken at 7 days after spraying. Scale bars: 1 cm.

## Discussion

Weeds compete with crops for limited resources of nutrition, space, light, and water and are thus undesirable plant species in agricultural production. In extreme cases when the weeds are left without any control, they may cause over 80% crop yield loss (Heap, 2014). Herbicides have been enthusiastically adopted by worldwide growers and have greatly accelerated the agricultural production efficiency and world crop production ever since they were developed. Based on mode of actions, herbicides could be further divided into different groups such as photosynthesis inhibitor, acetolactate synthase inhibitor, cellulose synthesis inhibitor (CBI), etc (Gianessi, 2013). Due to the natural mutation, herbicide tolerant weeds have become problematic after long time repetitive usage of specific leading herbicide (Delye et al., 2013; Heap, 2014). According to the international survey of herbicide resistant weeds (http://www.weedscience.org), 262 weed species (152 dicots and 110 monocots) have been reported to evolve herbicide resistance to 23 of the 26 known herbicide sites of action and to 167 different herbicides. Herbicide resistant weeds have been reported in 93 crops in 70 countries until 2019. Take 2,4-D for example, this well-known synthetic auxin has been commercialized ever since 1940s, and is one of the oldest and most widely used herbicides to control broadleaf weeds and woody plants in numerous small grain, fruit, and vegetable crops (Peterson et al., 2016). After over 70 years’ application, more than 40 weed species have already been reported to show resistance to 2,4-D according to the international survey of herbicide resistant weeds (http://www.weedscience.org). So finding novel herbicide is quite urgent to guarantee the crop production in order to feed the world ever increasing population.

CBIs are useful tool not only to understand cellulose biosynthesis, but also provide valuable resources for the commercial herbicide development. CBIs are the few herbicides which show less occurrence of weed tolerance (Heap, 2014). Most of the CBIs are discovered from chemical library screen and more than 10 CBIs have been identified so far (Tateno et al., 2016). Interestingly, the mutations at different CESAs, especially the primary cell wall related CESA 1, 3 and 6, have been found to cause reduced sensitivity to CBIs (Scheible et al., 2001; Desprez et al., 2002; Tateno et al., 2016; Hu et al., 2018). The CBIs and the reduced sensitivity mutants are valuable resources for the development of novel herbicides and for the breeding of herbicide tolerant crops which can be accomplished by gene editing technology (Hu et al., 2019).

Based on the effect of CBIs on CSC trafficking and the identified *CESA* mutants, it is reasonable to assume some CBIs may target the CESA directly. ES20 is a newly identified CBI that shares some characteristics with other known CBIs in terms of cellulose content reduction and ectopic accumulation of lignin and callose after treatment. The genetic and biochemistry evidences strongly support that ES20 targets CESA6 at the catalytic site (Huang et al., 2020). However, ES20 has a different mode of action than other three CBIs that we have tested based on a couple of observations. Firstly, most of the ES20 insensitive mutants are sensitive to other three CBIs whereas all three isoxaben insensitive mutants are sensitive to ES20. Secondly, all of the predicted ES20 binding site mutants are sensitive to the other three CBIs which indicate the binding pocket of ES20 is different than the other three tested CBIs.

Several ES20 insensitive mutants show cross tolerance to isoxaben, indaziflam and C17. For example, *es20r3* (CESA6^L935E^), *es20r4* (CESA6^D605N^) and *es20r5* (CESA6^S360N^) have reduced sensitivity to all four CBIs we have tested (Figure 3). The amino acids L935, D605 and S360 are important for CESA6 function because the mutants of *es20r3* (CESA6^L935E^), *es20r4* (CESA6^D605N^) and *es20r5* (CESA6^S360N^) have obvious root growth defects (Figure 3). The reduced sensitivities to CBIs caused by mutations in the same amino acids indicates these CBIs may share some common features in affecting cellulose synthesis although their exact target sites are different. It is possible that these amino acids are close to the target sites for these CBIs. This will remain as an open question because direct interaction between CESA and isoxaben, indaziflam and C17 needs further characterization. It is especially interesting for indaziflam because this CBI seems act differently than others at the cellular level. Indaziflam treatment causes increased CSC density at the PM, which is opposite to the effects of ES20, isoxaben, and C17 (Paredez et al., 2006; Brabham et al., 2014; Hu et al., 2016; Huang et al., 2020). It will be very interesting to investigate why the same mutations can lead to resistance to CBIs with different effects on CSC subcellular localization.

ES20 has synergistic inhibition effect on plant growth with other CBIs (Figure 6), which implies that it could be used together with other CBIs as herbicides to increase the weed control efficiency and to reduce the development of weed tolerance. ES20 could inhibit plant growth in soil condition, although a relative high dosage is needed (Figure 10). A future structure optimization will allow ES20 to be developed into a commercial herbicide with higher efficiency. Among 15 mutants that have reduced sensitivity to ES20, CESA6^E929K^ is the most efficient in tolerating the inhibitory effect of ES20. At the cellular level, PM-localized CESA6^E929K^ is not affected by ES20 treatment. ES20 tolerance caused by single amino acid change in CESA6 indicates that it is possible to create other ES20-tolerant crop species using gene editing technology. CRISPR-mediated gene editing has been suggested to obtain C17-tolerant plants (Hu et al., 2019) and it is possible to create ES20 tolerant plants as well with CRISPR technology. We also show that it is possible to create plants that have dual tolerance to ES20 and isoxaben. The double mutant of *ixr1-1*;*es20r1* show reduced sensitivity to the co-treatment of ES20 and isoxaben (Figure 9), indicating it is possible to create crops that are resistant to both ES20 and isoxaben. We did notice some slightly reduced root growth, smaller rosettes and shorter height in the double mutant of *ixr1-1*;*es20r1*, indicating spontaneous mutation at CESA3 and CESA6 further affects the normal function of CSC complex. The previous Quinoxyphen and isoxaben dual tolerant *CESA1* and *CESA3* double mutant *cesa1*^aegeus^/*cesa3*^ixr1-2^ also showed a far more pronounced dwarf phenotype than either of the single mutants (Harris et al., 2012). However, recently reported *CESA3^S983F^* and *CESA6^ixr2-1^* double mutant shows isoxaben and C17 dual tolerance but does not seem to have obvious growth phenotypes (Hu et al., 2019), which indicates different combination of mutated *CESAs* may affect the plant growth differently and it is still possible to obtain *CESA* dual drug tolerant plant without growth penalty. It is worth trying to create double amino acids mutations in *CESA6* for *irx2-1*;*es20r1* and test for the plants’ response to both ES20 and isoxaben and examine their growth phenotypes. Taken together, we have shown that ES20 has different mode of action than isoxaben, indaziflam and C17. ES20 has synergistic effect with isoxaben, indiaziflam and C17 in inhibiting plant growth. ES20 could be used as a potential spray herbicide and it is possible to create plants that can tolerate both ES20 and isoxaben by gene editing technologies.

## Materials and methods

### Plant material and growth conditions

To test the effect of ES20 on different plant species, Arabidopsis Col-0, tomato Micro-tom, soybean Williams 82, maize B73, and rice Nipponbare, perennial ryegrass Bright star and Kentucky bluegrass Brilliant were used. Dandelion seeds were collected from wild in West Lafayette, Indiana, USA. The seeds of Arabidopsis, dandelion, tomato, soybean, maize and rice were sterilized and sowed on half strength Murashige and Skoog medium (½ MS) with 0.8% agar at pH 5.8 with different concentrations of ES20. The plants were grown vertically under continuous light of 130 μmol m^-2^ s^-1^ intensity at 22 °C. Kentucky Bluegrass and Perennial Ryegrass seeds were directly grown in filter paper soaked in sterile water supplemented with DMSO (0.1%) or ES20 (50 μM) at 22 °C.

### Different CBI treatments

To test the sensitivity of *es20rs* and CESA binding site mutants to different CBIs, sterilized seeds were grown on ½ MS medium supplemented with indicated concentrations of CBIs. Equal volume of DMSO was used as a control. After 5 days of growth, the plates were scanned using Epson Perfection V550 scanner. The root length of plants was measured using ImageJ.

### Live cell imaging with spinning-disk confocal microscopy (SDCM)

SDCM was used to examine the localization of CSC at the PM. The seedlings of YFP-CESA6;*prc1-1* and YFP-CESA6^E929K^;*prc1-1* were grown on ½ MS medium for 5 days in vertical orientation. The seedlings were treated with DMSO or 6 μM ES20 for 30 min. Two thin strips of double-sided tape were placed on top of the glass slides about 2 cm apart from each other. 100 μl of ½ MS liquid growth media containing DMSO (0.1%) or 6 μM ES20 was applied to the glass slides with double-sided tape and then the seedlings were mounted in the liquid media carefully with tweezer. A 22 x 40 mm cover glass was placed on top of the double-sided tape for imaging. The images were taken from the 2nd or 3rd epidermal cells below the first obvious root hair initiation in the root elongation region. The SDCM that we used for imaging CSCs is a Yokogawa scanner unit CSU-X1-A1 mounted on an Olympus IX-83 microscope, equipped with a 100X 1.45–numerical aperture (NA) UPlanSApo oil objective (Olympus) and an Andor iXon Ultra 897BV EMCCD camera (Andor Technology). YFP fluorescence was excited with 515-nm laser line and emission collected through 542/27-nm filter.

### PM-localized CSC density analysis

To examine the effect of ES20 on PM-localized CSC density, images from SDCM were analyzed using ImageJ. The Freehand selection tool was used to choose region of interest (ROI) to avoid CSCs from Golgi. CSC particles from selected ROIs were detected on 8-bit images using the Find Maxima tool with the same noise threshold for all images. CSC particle density in ROIs was calculated by dividing the numbers of particles by the ROI area.

### ES20 spray test on soil grown plants

Arabidopsis Col-0 seedlings grown on ½ MS growth medium for 5 days were transferred to soil and covered with transparent plastic lid for 2 days. The plants were then sprayed with 50 mL sterile water supplemented with DMSO (0.5%) or ES20 (500 μM). The plants were imaged 7 days after spaying.

### Structure activity relationship analysis of ES20

To test the structure activity relationship of ES20, 11 ES20 analogs were ordered from Vitascreen (Champaign, IL, USA). Sterilized Arabidopsis Col-0 seeds were grown on ½ MS medium supplemented with 1 μM ES20 or different analogs. Equal volume of DMSO was used as a control. After 5 days of growth, the plates were scanned using Epson Perfection V550 scanner. The root length of plants was measured using ImageJ.

### Generation of transgenic Arabidopsis plants

YFP-*CESA6*^E929K^ construct was created as described previously (Huang et al., 2020). In brief, the genomic construct of CESA6 containing endogenous CESA6 promoter was cloned into modified binary vector pH7WGR2 with 35S promoter and RFP-tag removed. YFP-tag was inserted into the N-terminal region of the CESA6 start codon. The mutation E929K was introduced by site-directed mutagenesis. The verified plasmids were transformed into Col-0 or *CESA6* null mutant *prc1-1* (CS297) using *Agrobacterium tumefaciens* mediated floral dipping (Clough and Bent, 1998). The *prc1-1* seeds were obtained from the Arabidopsis Biological Resource Center (ABRC).

## Acknowledgments

We thank Dr. Yiwei Jiang for sharing the seeds of Kentucky Bluegrass and Perennial Ryegrass. We thank Dr. Cankui Zhang for sharing the rice seeds. We thank Dr. Christopher J. Staiger for allowing us to use the Spinning Disc Confocal Microscope for imaging CSC localization. The research is supported by Trask Trust Fund to C.Z and Purdue University provost’s startup fund to C. Z.

